# Microencapsulated *Vaccinium floribundum* Kunth extract promotes angiogenesis and attenuates inflammation in *in vitro* and *in vivo* models

**DOI:** 10.64898/2026.03.20.713210

**Authors:** Fabiana Antognoni, Irvin Tubon, Giulia Biondolillo, Luca Melotti, Roberta Di Lecce, Sherif M. Afifi, Roberta Salaroli, Gabriela Vaca, Cristian Vacacela Gomez, Goering Octavio Zambrano Cárdenas, Monica Forni, Augusta Zannoni, Chiara Bernardini

## Abstract

Natural products, especially polyphenol-rich medicinal plants, are increasingly investigated as multitarget therapeutics in both human and veterinary medicine for angiogenic regenerative properties and for inflammation based-diseases. Recent developments in natural product formulation, notably microencapsulation, have been shown to improve the stability, bioavailability, and controlled release of bioactive compounds. The integration of complementary *in vitro* and *in vivo* models is critical for evaluating both efficacy and translational potential. In this context, the present study assessed the phytochemical composition and biological activity of a microencapsulated Ecuadorian *Vaccinium floribundum* extract (VFM), using a combination of *in vitro* and *in vivo* approaches. VFM biochemical characterization identified 15 compounds, including flavonoids, procyanidins, dihydrochalcones, and phenolic acids, with chlorogenic acid and quercetin as the most abundant metabolites. Anthocyanins ideain and petunidin were also detected, confirming a rich bioactive profile. Primary porcine thoracic aortic endothelial cells (pAECs) were treated with VFM to assess cell viability and angiogenic potential and challenged with bacterial lipopolysaccharide (LPS) in the presence or absence of the extract. Anti-inflammatory effects were further evaluated *in vivo* using a carrageenan-induced mouse paw edema model. VFM enhanced endothelial cell viability, promoted capillary-like network and modulated early angiogenic signaling pathways. It mitigated LPS-induced endothelial dysfunction by reducing pro-inflammatory cytokines and oxidative stress markers. *In vivo*, paw edema assays confirmed its anti-inflammatory efficacy, with microencapsulation likely sustaining bioactive release. These findings support the traditional use of *Vaccinium floribundum* and highlight its potential for developing nutraceutical formulations targeting vascular and inflammatory disorders.

## 1 Introduction

In recent decades, there has been a substantial increase in scientific interest in natural products as therapeutic agents for both human and veterinary health. This trend is driven by the need to identify more safe and effective alternatives to synthetic drugs, particularly in the context of chronic and multifactorial diseases. Among natural source, medicinal plants have attracted particular attention due to their richness in bioactive metabolites, including phenolic compounds, flavonoids and terpenoids (1). These secondary metabolites are capable of modulating multiple molecular targets simultaneously. Such multitarget activity is especially relevant in complex biological processes that require fine regulatory balance. Inflammation and angiogenesis are closely interconnected biological processes that operate in a coordinated manner to regulate tissue homeostasis, repair, and adaptation to injury (2,3). Beyond their established roles in physiological contexts such as wound healing and embryonic development, the sustained activation of inflammatory and angiogenic pathways represents a hallmark of numerous pathological conditions, including chronic inflammatory diseases, cancer, and autoimmune disorders among others (4). Due to inflammation process occurring at both local and systemic levels, the integrated application of and in vivo models is essential for its comprehensive characterization. systems enable the detailed research of cellular responses and molecular signalling pathways under tightly controlled experimental conditions, whereas in vivo models recreate the complexity of the whole organism by incorporating systemic interactions and physiologically relevant immune dynamics. The complementary use of these approaches strengthens mechanistic interpretation and improves the translational assessment of potential therapeutic agents. Animal models have been fundamental for investigating normal physiology, understanding disease pathophysiology, and testing the safety and efficacy of new therapies in preclinical studies. This widespread usage is attributed to the genetic similarities between humans and various mammal species (5). Various studies have used different models to investigate inflammatory responses, highlighting the use of animals, especially rodents, to recreate acute inflammatory conditions. Moreover, these preclinical models not only facilitate the elucidation of underlying immunopathological mechanisms but also evaluate the pharmacodynamic efficacy and safety profiles of candidate anti-inflammatory agents (6). In this context, experimental models of acute inflammation are essential for understanding the underlying mechanisms and exploring potential therapeutic interventions. Large animal models play a crucial complementary role to small rodents by offering closer anatomical (7), physiological, and immunological similarity to humans. Among them, the pig model is particularly valuable in inflammation and angiogenesis research due to its comparable cardiovascular system, skin structure, wound healing response, and immune repertoire. Porcine models are widely used to study ischemia-induced angiogenesis, chronic inflammatory diseases, and biomaterial–tissue interactions. Among *in vitro* studies using the pig model, primary porcine Aortic Endothelial Cell cultures (pAEC) exhibit well-characterized morphological and molecular changes upon exposure to the pro-inflammatory lipopolysaccharide (LPS), including modulation of nitric oxide synthase expression, alteration of cell morphology, and induction of inflammatory signalling cascades, which closely mirror features of endotoxin-induced endothelial dysfunction observed *in vivo*. Importantly, this *in vitro* cell platform supports the evaluation of phytochemicals as protective agents against LPS-mediated damage, providing a controlled and reproducible experimental setting to assess their ability to preserve endothelial integrity and functionality under inflammatory conditions. The use of pAECs allows for the systematic investigation of plant-derived compounds in terms of their overall cytoprotective and anti-inflammatory potential, thereby offering valuable preliminary insights into their suitability for modulating endothelial responses to inflammatory insults (8–11). *Vaccinium floribundum* Kunth is a wild berry species of Ericaceae family growing in Ecuador’s Andean highlands, commonly known as mortiño (12). It is a semi-evergreen shrub, reaching 1.5 m in height, with lanceolate leaves that produce round, bluish-black berries with a pleasant flavour, mostly consumed in the local culinary tradition in jams, wine, and boiled drinks. Like most blueberries and cranberries, mortiño fruits are currently considered as “superfruits”, thanks to their high concentration of bioactive compounds mainly belonging to the class of phenolics, including polyphenols, flavonoids, anthocyanins, and proanthocynidins, and their well-recognized health-beneficial effects including antioxidant, antinflammatory, and antimicrobial ones (13,14). Therefore, it is crucial to ensure that bioactive compounds remain unchanged from their production until their consumption. To achieve this, the use of microencapsulation techniques, like spray drying, improves the stability, protection, and controlled release of bioactive compounds by the incorporation into a protective matrix, preserving the biochemical functionalities. This approach, thereby enhancing the efficacy and applicability of these extracts *in vivo* and *in vitro* trials (15). In this work the phytochemical composition and the biological potential of a microencapsulated extract of *Vaccinium floribundum* berries was evaluated, with the aim of providing a scientific support to the anti-inflammatory and the angiogenic properties of this plant, using *in vitro* and *in vivo* models.

## 2 Materials and methods

### 2.1 Chemical and Reagents

Cell culture plates and glass slides were obtained from Corning Life Sciences (NC 27712 USA).Human Endothelial Serum-Free Medium (hESFM), heat-inactivated fetal bovine serum (FBS), antibiotic-antimycotic, Dulbecco’s phosphate-buffered saline (DPBS), phosphate-buffered saline (PBS), and Geltrex™ LDEV-Free Reduced Growth Factor Basement Membrane Matrix were purchased from Gibco-Life Technologies (Carlsbad CA, USA). Trypsin–EDTA solution 1X, Dimethyl sulphoxide (DMSO), lipopolysaccharide (LPS) (E. coli 055:B5), *In vitro* Toxicology Assay Kit and Cell Growth Determination Kit MTT based and Maltodextrin (DE 10–12) was used as wall material for microencapsulation were purchased from Merck-Sigma-Aldrich (St. Louis, MO, USA). TRIzol reagent was purchased from Thermo Fisher Scientific (Waltham, MA, USA). RNA isolation was performed with a NucleoSpin RNA II kit (MACHEREY NAGEL GmbH & Co. KG, Düren, Germany), and an iScript cDNA Synthesis Kit and iTaq Universal SYBR Green Supermix were used for cDNA synthesis and qPCR analysis, respectively (Bio-Rad Laboratories Inc., Hercules, CA, USA). The absence of mycoplasma contamination was periodically verified using the EZ-PCR Mycoplasma Detection Kit (Biological Industries, Kibbutz Beit-Haemek, Israel). Chemical standards (chlorogenic, caffeic, *p-*coumaric, ferulic, protocatechuic acid, quercetin, quercetin-3-*O*-galactoside, quercetin-3-*O*-glucoside, quercetin-3-*O*-rhamnoside, quercetin-3-O-rutinoside, kaempferol, kaempferol-7-*O-*glucoside, cyanidin-3-*O*-galactoside chloride, catechin, epicatechin, procyanidin B1, procyanidin B2, procyanidin C1, phloretin, and phlorizin), all >95% pure, were purchased from Extrasynthese (Genay Cedex, France). Stock solutions of the analytes were prepared at 1 mg/mL in MeOH and stored at - 20°C until use. Working standard solutions were prepared daily from stock solutions by dilution with the corresponding mobile phase in its initial composition for phenolic acid and flavonoid analysis, respectively. HPLC-grade methanol (MeOH), acetronitrile, and water, formic acid, fluorescein, K_2_HPO_4_, 2,2’-azobis(2-amidinopropane) dihydrochloride (AAPH), 2,2-diphenyl-1-picrylhydrazyl (DPPH), and Trolox (Tx), all pure for analysis, were purchased from Merck Reagents (Milan, Italy).

### 2.2 Collection of plant material

The fruits of *Vaccinium floribundum* Kunth (mortiño), were collected, after obtaining authorization for non-commercial collection from the Ministry of Environment, Water and Ecological Transition (No. MAATE-ARSFC-2022-2801), in the province of Tungurahua, specifically in the parish of Pilahuín, at a latitude (Y): −78.7277812; longitude (X): −1.299677 and altitude: 3,054 meters above sea level. This process was carried out using simple random sampling to ensure the quality of the samples collected. The fruits were collected when they were in good condition, without damage from insects, animals, or environmental factors such as rain, sun, or wind.The plant material was identified by the herbarium of the Escuela Superior Politécnica de Chimborazo (No-ESPOCH-HERB-035) and subsequently stored at the Pharmaceutical Compounding Laboratory of the Escuela Superior Politécnica de Chimborazo.

### 2.3 Preparation of plant material

Sodium hypochlorite solution at 9% was used to disinfect the fruits and subsequentially washed with abundant water to eliminate excess chlorine. Then a convection dehydrator (Memmert, USA) was used to dry the material at 40 °C for 48 to 72 hours. Finally, the dried material was crushed using a blender until fine powder was obtained and stored in ziplock bags (16).

### 2.4 Extract preparation

The maceration method was used to obtain ethanolic extract from dry and crushed fruits. Thus, dried fruits with 50:50 ethanol: water ratio were placed in a glass container with a 1:10 ratio (w/v) and left to stand in the absence of light for 8 days, shaking occasionally to distribute the plant material in the solvent. After this time, the product was filtered, and the solvent was removed by rotary vacuum evaporation (Bucchi, Switzerland). Finally, the mortiño ethanolic fruit extract (VFE) was stored in sterile plastic tubes under refrigeration at 4°C until use (17).

### 2.5 Microencapsulation Process

The microencapsulation of plants extracts was performed using maltodextrin (10–20 DE) as the carrier agent. Extracts with a total solids content of 30% (equivalent to 20 g solids) were prepared at a 20:80 solid extract-to-maltodextrin ratio. For each extract, 280 mL of distilled water was mixed with the anthocyanin solution and combined with maltodextrin. The microencapsulation process using a laboratory-scale mini spray dryer Büchi B-290 (Flawil, St. Gallen, Switzerland). The solutions were fed into the spray dryer under controlled conditions: an inlet air temperature of 140 ± 2 °C, an outlet air temperature of 80 ± 2 ◦C, an internal pressure of −50 mbar, an atomizing air flow of 400–600 L/h, and a drying air flow rate of 60 m^3^/h. During the spray drying process, each solution was atomized within the drying chamber, forming solid microspheres encapsulated in maltodextrin-based polymers. The resulting microspheres of the mortiño ethanolic fruit extract microencapsulated (VFM) were collected and stored in HDPE-aluminum bags at room temperature of 15–25 °C (18).

### 2.6 Encapsulation yield

Encapsulation yield after spray drying was calculated as a percentage as a ratio between the total microcapsules powders weight and the weight of the carrier agents using the following equation:

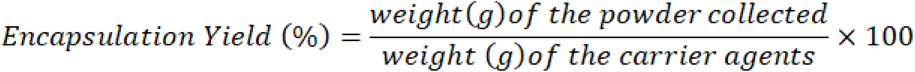

### 2.7 Characterization by FTIR

Fourier transform infrared (FTIR) spectroscopy was employed to identify the functional groups in the microcapsules. he spectra were obtained using a Perkin Elmer Spec-trum two FTIR L1600312 spectrometer (Perkin Elmer, Tokyo, Japan) with the ATR method, covering a wave number range of 4000–500 cm⁻^1^ at a resolution of 4 cm^−1^ across 36 scans (19).

### 2.8 HPLC-DAD analysis of phenolic acids and flavonoids in VFM

The identification and quantification of phenolic acids was performed according to Mattila and Kumpulainen (20) with modifications (21), using a Jasco (Tokyo, Japan) HPLC-DAD 4000 series system composed of a PU-4180 pump, an MD-4015 photodiode array detector, a XX oven and an AS-4050 autosampler. The stationary phase was an Agilent (Santa Clara, CS, USA) Zorbax Eclipse Plus C18 reverse-phase column (4.6 x 100 mm, 3.5 µm) equipped with a Zorbax Eclipse Plus C18 precolumn. The mobile phase was composed by 4.5% of formic acid (FA, solution A) and acetonitrile (ACN, solution B) and the elution was carried out at a flow rate of 0.7 mL/min and a temperature of 24°C, as follows: isocratic elution 97% A, 0 – 6 min; linear gradient from 97% A to 85% A, 6 – 17 min; linear gradient from 85% A to 82.8% A; 17 – 27 min; linear gradient from 82.8% A to 50% A, 27 – 40 min; linear gradient from 50% A to 97% A, 40 – 43 min; isocratic elution 97% A, 43 – 50 min. The injection volume for both standards and samples was 20 µL and analyte detection was performed by monitoring at 329, 287, 280, and 254 nm. Quantifications were carried out using pure standards and building calibration curves in the 0.3 - 80 ppm range for all considered analytes. The identification and quantification of flavonoids was performed according to the method by Wojdyło et al. (22) with modifications (9), using the same chromatographic system and column as described above. The mobile phase was composed by 50 mM phosphoric acid, pH 2.5 (solution A) and acetonitrile (solution B) and the elution was carried out at a flow rate of 0.5 mL/min and a temperature of 24°C as follows: isocratic elution 97% A, 0 – 1 min; linear gradient from 97%A to 60%A, 1 – 31 min; linear gradient from 60% A to 36% A; 31 – 35 min; linear gradient from 36% A to 97% A, 35 – 37 min; isocratic elution 97% A, 37 – 47 min. The injection volume for both standards and samples was 20 µL and analyte detection was performed by monitoring at 360 and 280 nm. Quantifications were performed using pure standards and building a calibration curve in the range 0.3-80 ppm for all considered analytes.

### 2.9 HPLC-MS analysis of anthocyanins in VFM

The chromatographic system was an Agilent 1260 Infinity coupled to Agilent Ultivo 6465B triple quadrupole LC-MS (Agilent Technologies Inc., Santa Clara, California, USA). The chromatographic separations were achieved by a Zorbax SB-C18 (4.6×150 mm, 3.5 μm) using a mobile phase composed of 0.1 % FA in ACN (Component A) and 0.1 % FA in water (Component B) flowing at 0.40 mL/min. The gradient elution program was optimized as below: 0 min, 5% A; 3 min, 10% B; 35 min, 11% B; 40 min, 20% B; 50 min, 30% B; 55 min, 5% B and held constant for 5 additional minutes for column reconditioning. The injection volume was 10 μL and injections were carried out through the autosampler integrated into the Agilent system.

The data were carried out in multiple reaction monitoring (MRM) scan mode, using electrospray ionization source in positive mode. The main parameters for the mass spectrometer were as follows: capillary voltage: 4.0 kV; gas temperature: 300°C; gas flow: 13 L/min; nebuliser: 30 psi; sheath gas temperature: 300°C; sheath gas flow: 10 L/min. The specific, analyte-dependent parameters are detailed in Table 1.

**Table 1.** Optimised MRM MS parameters.

| Compound | Ion Mode | Precursor ion (*m/z*) | Product ion (*m/z*) | Collision energy (V) |
| --- | --- | --- | --- | --- |
| Cyanidin-3-*O*-galactoside (ideain) | + | 449 | 287 | 25 |
| Petunidin-3-*O*-glucoside | + | 479 | 317 | 25 |
| Peonidin-3-*O*-glucoside | + | 463 | 301 | 25 |
| Delphinidin | + | 303 | 229 | 40 |
| Cyanidin | + | 287 | 213 | 40 |

### 2.10 *In vitro* antioxidant activity assays

The *in vitro* antioxidant capacity of the microencapsulated extract was evaluated through the DPPH (2,2-diphenyl-1-picrylhydrazide) and the *Oxygen Radical Absorbance Capacity* (ORAC) assays as previously described (8).

### 2.11 Cell culture and treatment

Primary porcine aortic endothelial cells (pAECs) were isolated, expanded, and characterized as previously described (23). After isolation and expansion, cells were cryopreserved until further required. For experimental procedures, pAECs were seeded and routinely cultured in T25 or T75 flasks (2 × 10⁴ cells/cm²) in Human Endothelial Serum-Free Medium (hESFM) supplemented with 5% Fetal Bovine Serum (FBS) and 1% antibiotic–antimycotic solution. Cells were maintained at 38.5 °C in a humidified atmosphere containing 5% CO₂.

Cell cultures were routinely checked for mycoplasma contamination. At approximately 70–80% confluence, cells were exposed to the experimental treatments.

VFM extract was tested at increasing concentrations (1 ng/mL, 10 ng/mL, 100 ng/mL, 1 μg/mL, 10 μg/mL, 100 μg/mL, and 1 mg/mL) for cell viability assessment. Based on viability results, selected concentrations of VFM (0.1 and 1 mg/mL) were used for subsequent experiments, either alone or in combination with lipopolysaccharide (LPS; 10 μg/mL). For gene expression analyses (qPCR), confluent pAECs (approximately 1 × 10^5^ cells/well) were treated with the highest and most effective concentration based on viability test of VFM (1 mg/mL) in the absence or presence of LPS (25 µg/mL) for 5 h.

For the capillary-like tube formation assay, cells were seeded onto GelTrex™ -coated chambers and exposed to VFM (0.1 or 1 mg/mL), and tube formation was evaluated after 5 h.

Untreated cells cultured under identical conditions served as controls in all experiments.

### 2.12 Cell viability

Cell viability was evaluated using an MTT-based assay Briefly: sixth-passage pAECs were seeded in 96-well culture plates at a density of 2×10^4^ cells/well and incubated for 24 h. Afterwards, the culture medium was then replaced with hESFM supplemented with 5% FBS containing increasing concentrations of VFM or LPS, alone or in combination and cells were incubated for an additional 24h at 38,5°C and 5% CO_2_.

After treatment, cells were incubated with MTT solution (0.5 mg/mL) for 3 h at 38,5°C and 5% CO_2_. Subsequently, 0.1 mL of MTT solubilisation solution was added to each well to dissolve the formazan crystals.

Absorbance was measured at 570 nm using Infinite® F50/Robotic microplate reader (TECAN Life Sciences, Männedorf, Switzerland). For blank subtraction wells with only medium were used.

### 2.13 Quantitative Real Time PCR

RNA extraction was performed from pAECs seeded in a 24-well plate exposed to VFM (1mg/mL) alone or in combination with LPS (25 μg/mL), for 5 h or 24 h by using the TRIzol reagent and NucleoSpin RNA II kit, according to the manufacturer’s instructions.

After medium removal, 500 µL of TRIzol reagent were added to each well, and cells were lysed for 3 minutes at room temperature. Lysates were transferred into microcentrifuge tubes, mixed with 200 µL of chloroform, incubated for 10 minutes, and centrifuged (12000 xg for 15 min). The aqueous phase was recovered, mixed with an equal volume of 70% ethanol, and the resulting solution was applied to the NucleoSpin RNA Column for purification.

After spectrophotometric quantification, total RNA (250 ng) was reverse-transcribed into cDNA using the iScript cDNA Synthesis Kit (Bio-Rad) in a final volume of 20 μL.

All amplification reactions were performed in 20 μL and analyzed in duplicates (10 μL/well). The multiplex PCR contained the following: 10 μL of iTaqMan Probes Supermix (Bio-Rad), 1 μL of forward and reverse primers (5 μM each) of each reference gene, 0.8 μL of iTaqMan probes (5 μM) of each reference gene, 2 μL of cDNA, and 2.6 μL of water. The following temperature profiling was used: initial denaturation at 95°C for 30 seconds followed by 40 cycles of 95°C for 5 seconds and 60°C for 30 seconds.The SYBR Green reaction contained the following: 10 μL of iQ SYBR Green Supermix (Bio-Rad), 0.8 μL of forward and reverse primers (5 μM each) for each target gene, 2 μL of cDNA, and 7.2 μL of water. The real-time program included an initial denaturation period of 1.5 min at 95°C, 40 cycles at 95°C for 15 s, and 60°C for 30 s, followed by a melting step with ramping from 55°C to 95°C at a rate of 0.5°C/10 s. To evaluate gene expression profiles, quantitative real-time PCR (qPCR) was performed using a CFX96 thermal cycler (Bio-Rad). Expression levels of target genes (VEGF; FLT1; FLK1, HemeOoxygenase-1, HO-1 Interleukin-6, IL-6; Interleukin-8, IL-8;) were analyzed using SYBR Green chemistry (8,9,24,25), while reference genes (Glyceraldehyde-3-phosphate dehydrogenase, GAPDH; β-Actin, β-ACT) were assessed using TaqMan probes in multiplex reactions (26). Primer sequences, expected PCR product lengths, and NCBI accession numbers are shown in Table 2. Relative gene expression levels were normalized to the geometric mean of the two reference genes. The relative mRNA expression of the tested genes was calculated using the 2^-ΔΔCT^ method (27) and referred to pAECs cultured under the standard condition (control).

**Table 2.**
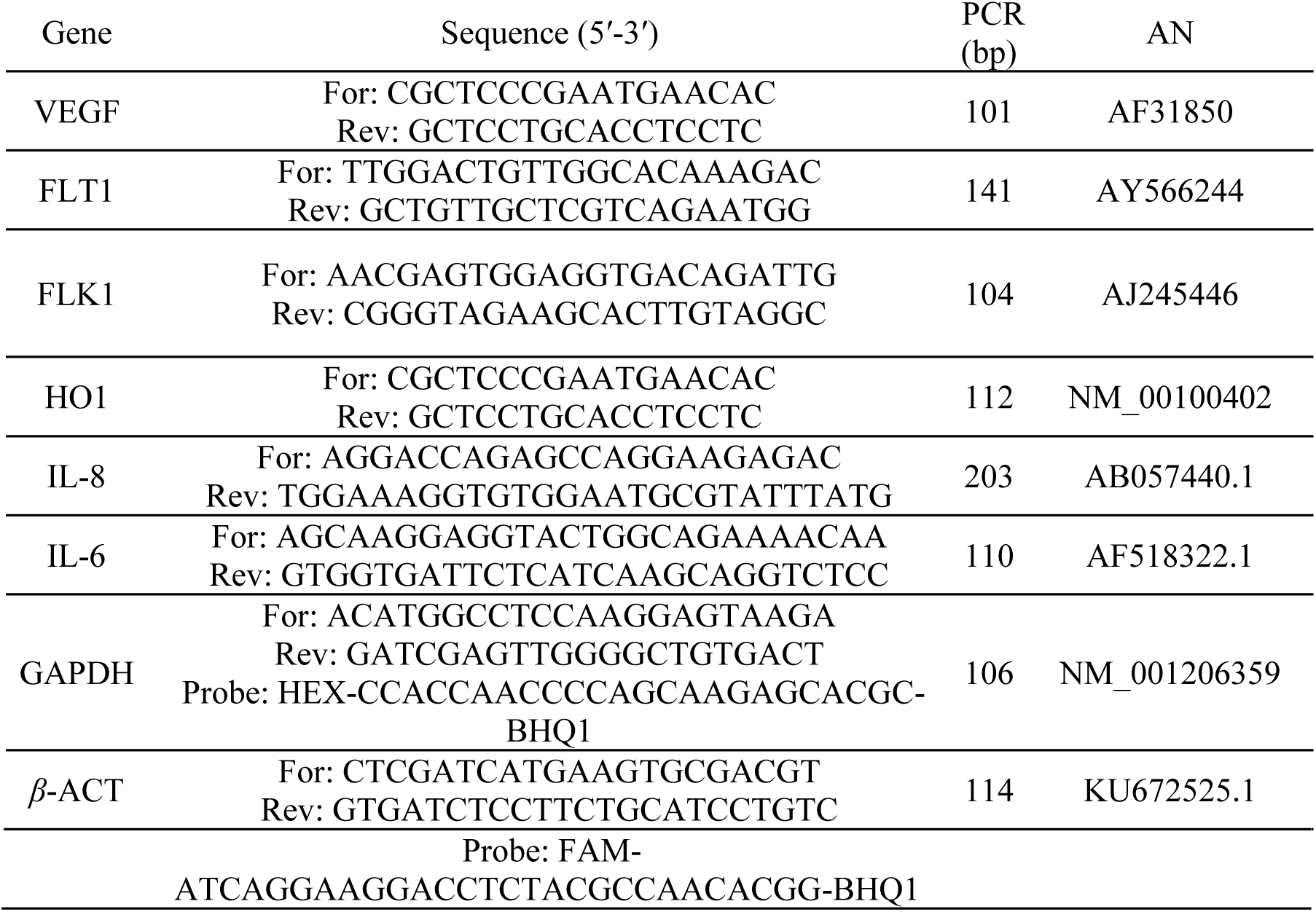
Primer sequence used for quantitative real time PCR analysis. AN = Accession Number (NCBI)

### 2.14 *In vitro* Capillary-Like Tube Formation Assay

The ability of pAECs to form capillary-like structures was evaluated using a tube formation assay on a three-dimensional extracellular matrix. Experiments were conducted in 8-well chamber slides coated with GelTrex™ Reduced Growth Factor Basement Membrane Matrix (Gibco, Thermo Fisher Scientific), as previously described (24). GelTrex™ was added to each well and allowed to polymerize for 30 min at 38.5 °C in a humidified atmosphere containing 5% CO₂.After polymerization, pAECs were seeded onto the matrix at a density of 1 × 10⁵ cells/well. Cells were cultured in hESFM without FBS or antibiotics, to avoid interference with angiogenic responses. Cells were treated with VFM (0.1 or 1 mg/mL) and incubated under standard culture conditions.Images were acquired using a digital camera installed on a Nikon contrast phase microscope (Nikon, Yokohama, Japan). Quantitative analysis of tubular structures was performed using ImageJ 64 software with the Angiogenesis Analyzer plugin.

### 2.15 Animals

Male Balb/c mice (27.5±2.5 g) were obtained from the Bioterium of Science Department of the Escuela Superior Politécnica de Chimborazo. They were group-housed in cages at temperature (24 ± 1 °C), constant humidity (55 ± 5 %), and light–dark cycle (on at 7:00 a.m. and off at 7:00 p.m.). All animals were allowed *ad libitum* food and water according to the experimental protocols. Studies were performed in accordance with the NIH Guide for the Care and Use of Laboratory Animals, and the protocol was approved by the Ethics Committee of the Escuela Superior Politécnica de Chimborazo (ESPOCH-CIBE-2022-0037).

### 2.16 Carrageenan-induced mouse paw edema

Thirty mice were randomly divided into 5 groups (5 mice per group): blank control, negative control (Carrageenan 1%), positive control (Carrageenan 1% + diclofenac suspension at 10 mg/kg), and experimental groups (Carrageenan 1% + VFM at 75, 150 and 300 mg/kg). The vehicle used to solubilize sodium diclofenac and the extracts administered orally was 0.5% w/v carboxymethylcellulose. Acute inflammation was produced by sub-plantar injection (0.25 μL) of 1% freshly prepared suspension of carrageenan into the right hind paw. The paw volume was measured with a plethysmometer to determine the difference between the right and left paws at 1, 2, 3, 4, 5, and 6 h after carrageenan injection. The percentage inhibition of edema was calculated by comparison of the control group and experimental treatments as previously described (28). At the end of the experiment, animals were euthanized using an intraperitoneal overdose of anesthetic (ketamine/xylazine, Agrovet-Market). This procedure was performed in accordance with international recommendations for the use and care of laboratory animals and to minimize pain, stress, and suffering.

## 3 Statistical Analysis

Data were analysed by a one-way analysis of variance (one way ANOVA) followed by the “post-hoc” Tukey comparison Test or by Student’s *t*-test, as appropriate. Differences of at least *p* ≤ 0.05 were considered significant. Statistical analysis was carried out using GraphPad Prism 7 software.

## 4 Results

### 4.1 Phytochemical composition of VFM

#### 4.1.1 Yield

Through the microencapsulation process, a total of 19.07 g of microencapsulated material was obtained, yielding 75.97% as shown in table 3.

**Table 3.** Percentage yield of Mortiño Fruit extract (VFM) microencapsulated.

| Microencapsulated | Concentration<br>(ethanol: water) | Yield (%) |
| --- | --- | --- |
| Mortiño Fruit extract (VFM) | 50:50 | 75.97 |

#### 4.1.2 Structural Characterization by FTIR

The FTIR spectra of extracts, before and after microencapsulation, revealed characteristic modifications that confirm the successful entrapment of polyphenolic compounds within the maltodextrin matrix. The most evident change was the attenuation of the broad O–H stretching band (3200–3600 cm−1) and the reduction in C=O stretching intensity (1700–1750 cm−1) in the encapsulated sample. These shifts indicate hydrogen bonding interactions between polyphenols hydroxyl/carbonyl groups and maltodextrins, or steric shielding of reactive groups within the carbohydrate network (Figure 1).

**Figure 1:**
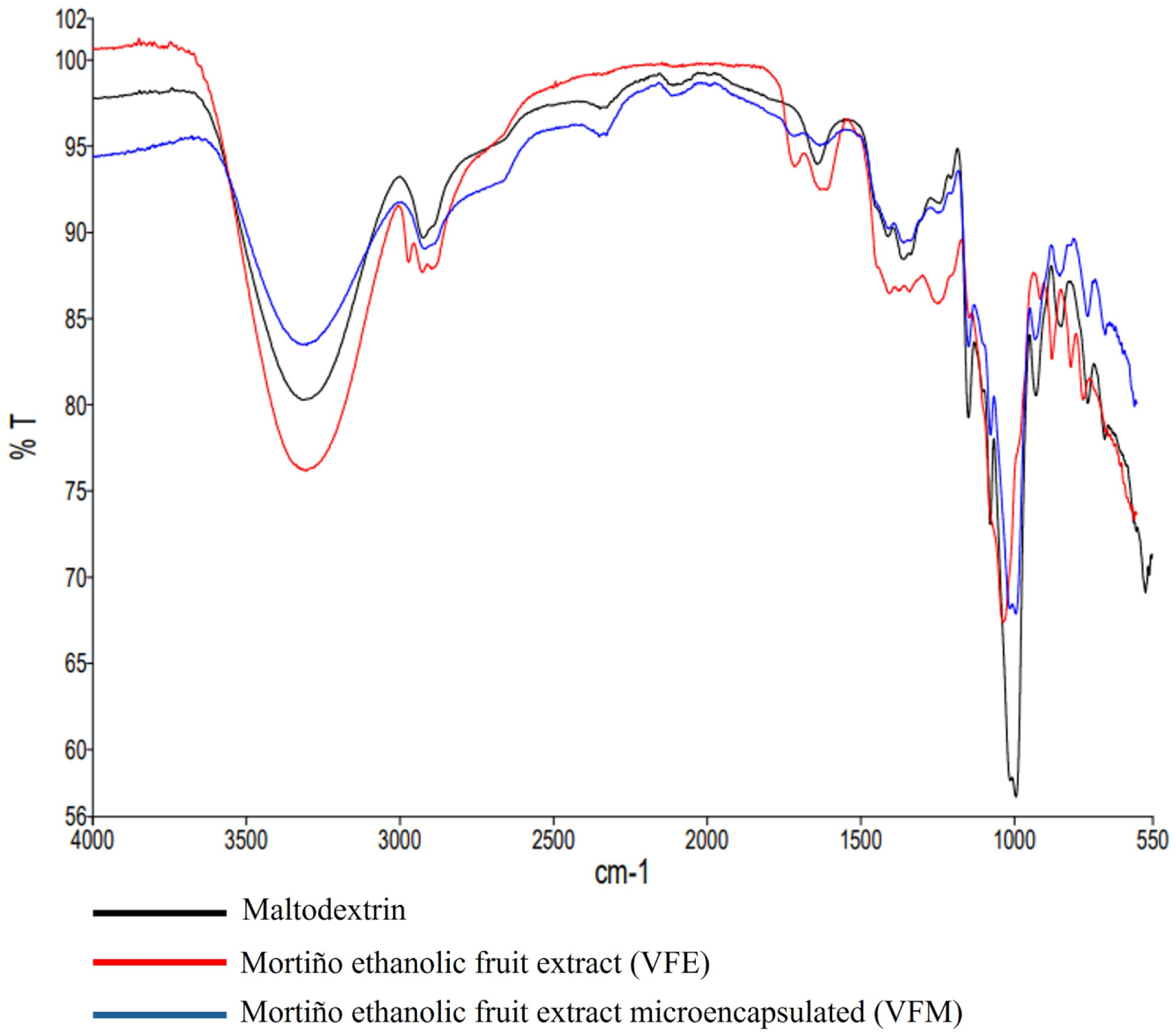
FTIR spectra before and after microencapsulation of mortiño fruit extract.

A total of 15 compounds were identified and quantified using HPLC–DAD analysis of VFM, nine of which were flavonoids (Table 4). A diverse profile of secondary metabolites, specifically within the classes of procyanidins, flavonols, and dihydrochalcones, was found (Table 3). Procyanidin C1 (2.03 ppm) and procyanidin B2 (0.23 ppm) were considerably less abundant than procyanidin B1 (5.94 ppm). Quercetin (5.29 ppm) was the most common aglycone in the flavonol category while its glycoside counterpart hyperoside reached 2.68 ppm. In contrast, the glycosylated form kaempferol-7-*O*-glucoside (3.30 ppm) surpassed the aglycone kaempferol (1.82 ppm) for kaempferol derivatives. Interestingly, similar quantities of the dihydrochalcone phloretin and its 2’-glucoside phloridzin were detected (1.25 ppm). Lastly, the flavan-3-ol monomer epicatechin was detected at 1.13 ppm. Regarding the quantification of phenolic acids, chlorogenic acid was the most prevalent metabolite in VFM, with a concentration of 13.21 ppm. Protocatechuic acid, a hydroxybenzoic acid, followed next at 0.37 ppm. The concentrations of hydroxycinnamic acids, i.e., caffeic, ferulic, and *p*-coumaric acids were considerably lower and nearly equal at 0.27 ppm. LC-MS analysis of anthocyanins in VFM revealed the presence of ideain (cyanidine-3-*O*-galactoside) and petunidin, while cyanidin and peonidin were not detected (below the limit of quantification) (Table 4).

**Table 4.** Quantified phenolics and antioxidant activity in VFM through HPLC-DAD and HPLC/MS. The concentration of identified compounds is expressed as ppm ±SD; the antioxidant capacity is expressed as nmol Trolox Equivalents/mg dry weight.

| Class of compounds/Assay | Compound/assay | Concentration or activity |
| --- | --- | --- |
| Phenolic acids | Protocatechuic acid | $0.37 \pm 0.17$ |
| | Caffeic acid | $0.27 \pm 0.07$ |
| | Chlorogenic acid | $13.21 \pm 2.01$ |
| | Ferulic acid | $0.27 \pm 0.01$ |
| | *p*-Coumaric acid | $0.27 \pm 0.02$ |
| Flavonoids | Quercetin | $5.29 \pm 0.10$ |
| | Quercetin-3-*O*-galactoside | $2.68 \pm 0.36$ |
| | Kaempferol | $1.82 \pm 0.07$ |
| | Kaempferol-7-*O*-glucoside | $3.30 \pm 0.10$ |
| | Phloretin | $1.25 \pm 0.25$ |
| | Phloridzin | $1.28 \pm 0.25$ |
| | Epicatechin | $1.13 \pm 0.17$ |
| | Procyanidin B1 | $5.94 \pm 2.70$ |
| | Procyanidin B2 | $0.23 \pm 0.01$ |
| | Procyanidin C1 | $3.65 \pm 0.67$ |
| Anthocyanins | Cyanidin | n.d. |
| | Cyanidin-3-*O*-galactoside | $2.58 \pm 0.06$ |
| | Petunidin | $0.80 \pm 0.01$ |
|  | Peonidin | n.d. |
| Antioxidant activity | DPPH | $38.86 \pm 0.41$ |
| | ORAC | $48.89 \pm 1.73$ |

### 4.2 *In vitro* model

#### 4.2.1 Angiogenesis

##### 4.2.1.1 Effects of VFM on Endothelial Cell Viability

Treatment of pAECs with VFM for 24 h did not negatively affect cell viability at any tested concentration. Nevertheless, exposure to VFM at 1 mg/mL (the highest concentration) significantly increased cell viability compared with the control condition (p < 0.05) (Figure 2).

**Figure 2:**
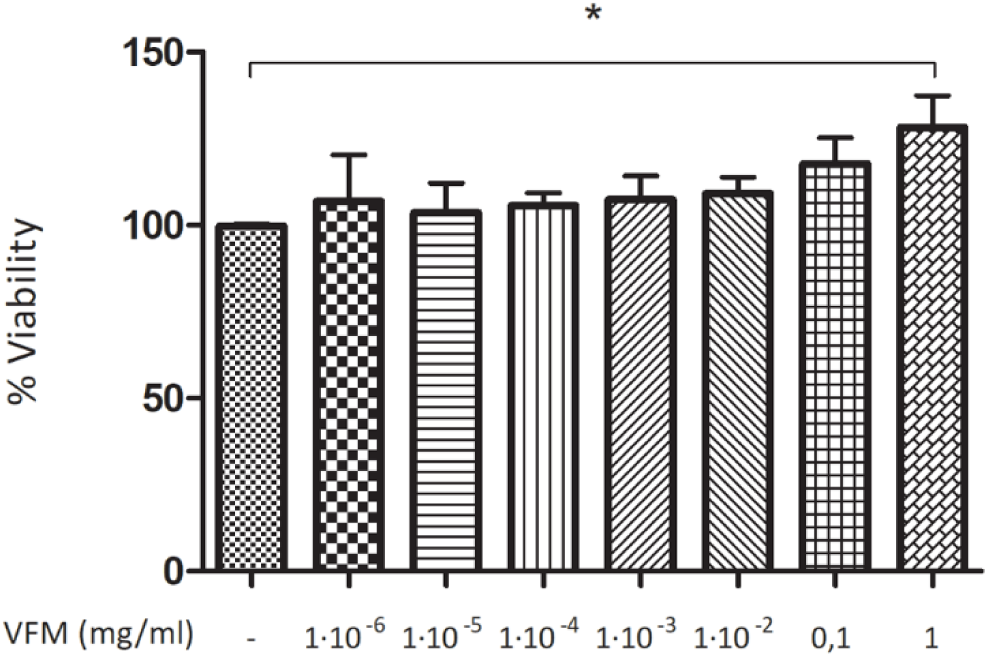
Effects of VFM on cell viability assessed by the MTT assay. pAECs were treated with increasing concentrations of VFM for 24 h. Statistical analysis was performed using Student’s t-test comparing each treated group versus the untreated control (CTR). Data are presented as mean ± SEM. n=3. CTR (-), untreated control; VFM, cells treated with the Vaccinium floribundum Kunth microencapsulated extract. * p < 0.05.

##### 4.2.1.2 Effects of VFM on Angiogenesis-Related Gene Expression

The expression of angiogenesis-related genes, including VEGF, its receptors FLK-1 (VEGFR-2) and FLT-1 (VEGFR-1), and HO-1, was evaluated at 5 h and 24 h following treatment with the phytoextract.

At 5 h (Figure 3A), a statistically significant effect was observed exclusively for FLT-1 expression showing an up-regulation of its gene expression in the presence ofVFM at 0.1 mg/mL, whereas no significant difference was observed for other genes.

**Figure 3:**
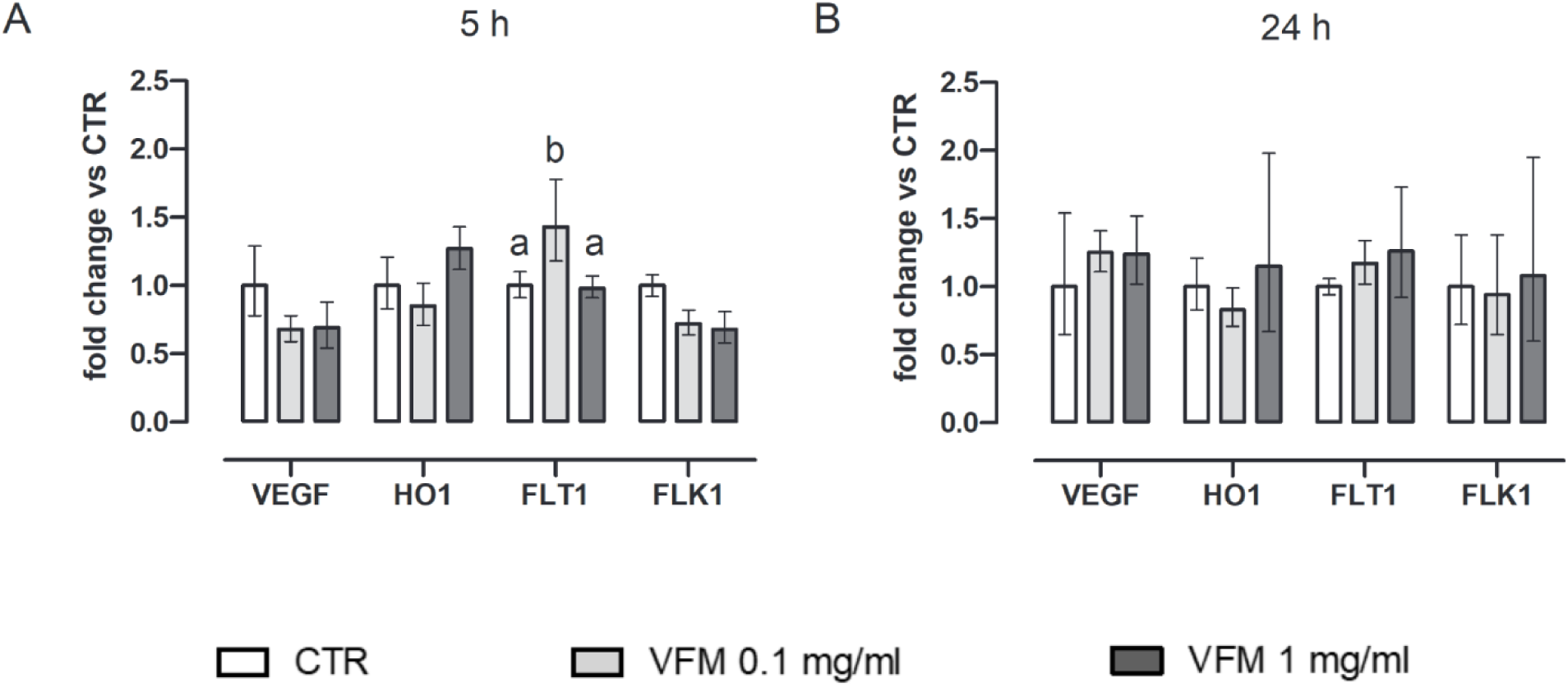
Effect of VFM treatment on angiogenesis-related gene expression. Expression of VEGF, HO-1, FLT-1 (VEGFR-1), and FLK-1 (VEGFR-2) was evaluated by qPCR after 5 h (A) and 24 h (B) of treatment with VFM (0.1 mg/mL and 1 mg/mL). Gene expression levels are reported as Fold Change (2-ΔΔCt) in respect to the control cells. Error bar represents the range of relative gene expression (n=3). Statistical analysis was performed using one-way ANOVA, followed by appropriate post-hoc multiple comparison tests. Different letters indicate statistically significant differences among groups (p < 0.05). CTR, untreated control; VFM, *Vaccinium floribundum* Kunth microencapsulated extract.

After 24 hours of treatment, no differences were observed for the mRNA levels of studied genes (Figure 3B).

##### 4.2.1.3 *In vitro* Capillary-Like Tube Formation Assay

To investigate the pro-angiogenic potential of VFM, an *in vitro* capillary-like tube formation assay was performed on pAECs. Representative images obtained after 5 h of treatment with 1 mg/mL VFM showed well-defined capillary-like networks composed of interconnected junctions, segments, and meshes (Figure 4). All morphometric parameters were evaluated for untreated control cells (CTR) and for pAECs treated with VFM at concentrations of 0.1 mg/mL and 1 mg/mL, quantitative analysis revealed statistically significant differences between control and treated samples for all evaluated parameters (Figure 5).

**Figure 4:**
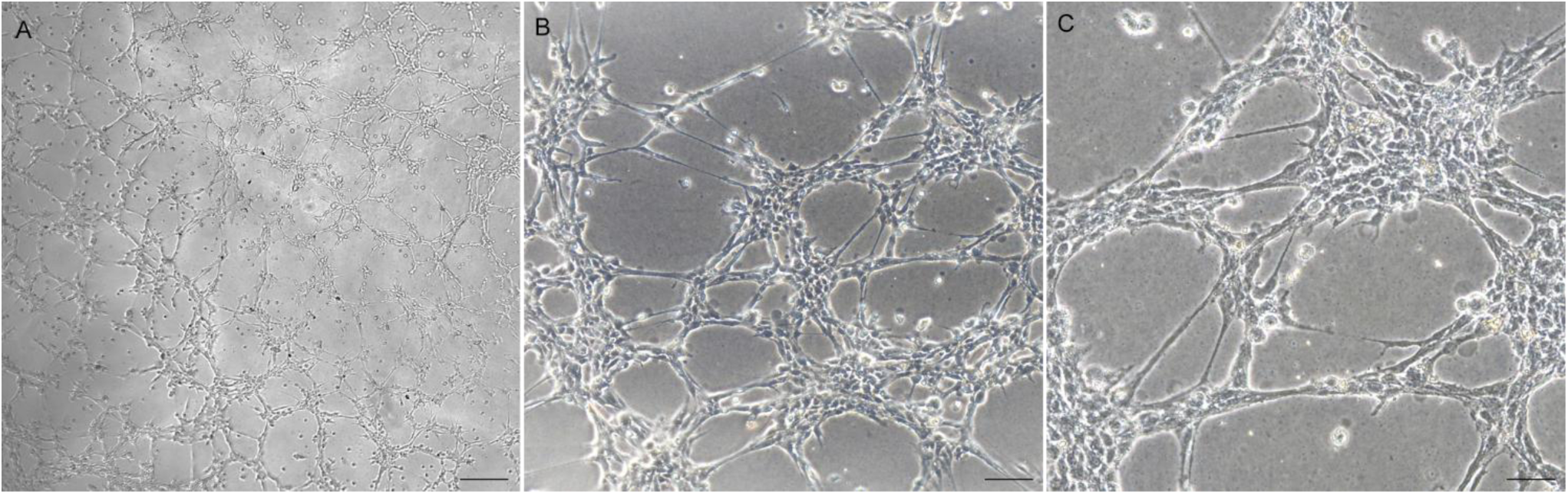
Representative phase-contrast images of tubular structure formation in pAECs treated with VFM at 1 mg/mL (VFM 1) after 5 h of incubation. (a) scalebar = 500 µm., (b) scalebar = 100 µm (c) scalebar = 50 µm

**Figure 5:**
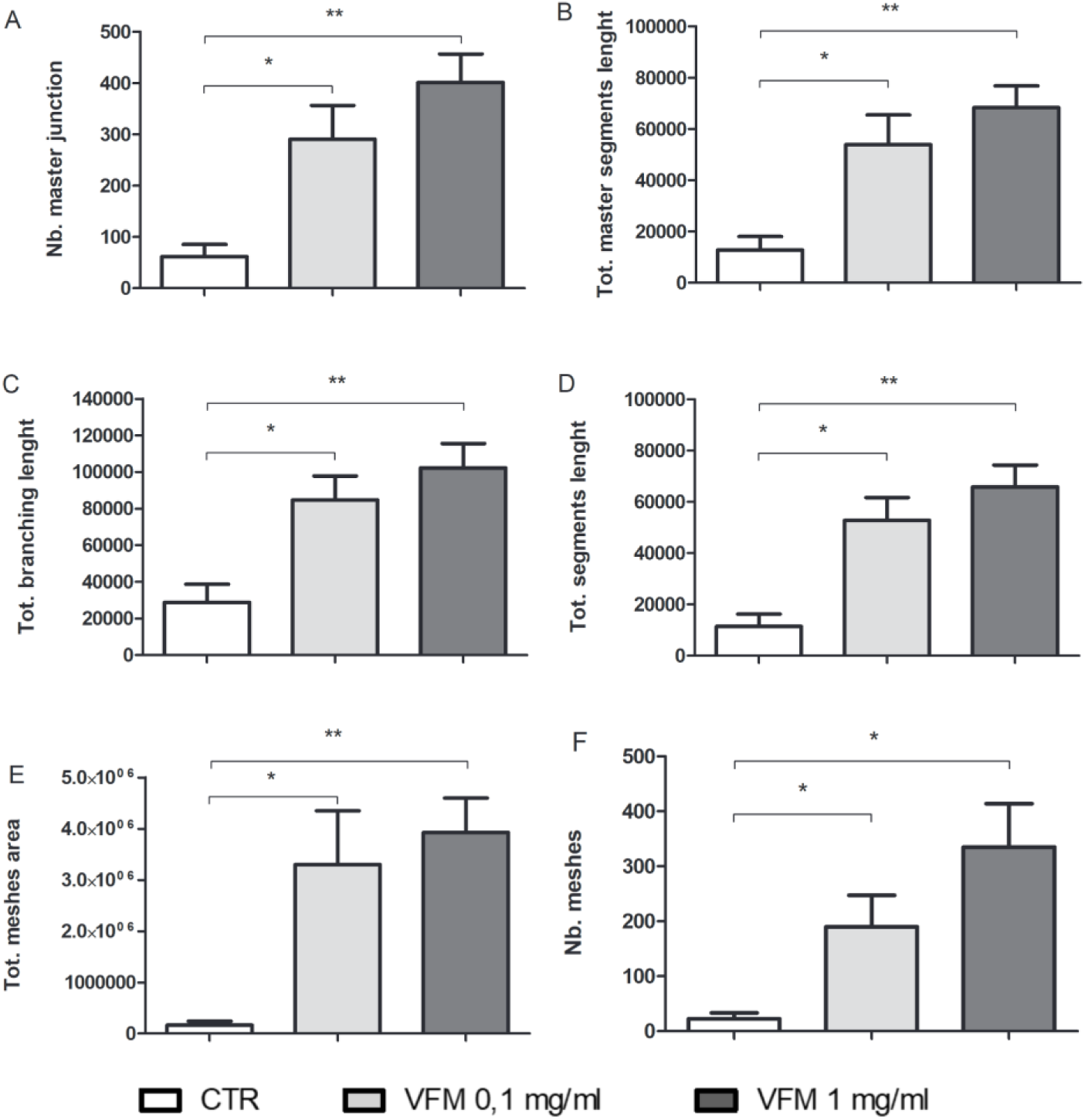
*In vitro* capillary-like tube formation assay following VFM treatment. Morphometric analysis of capillary-like networks formed by pAECs treated with VFM (0.1 mg/mL and 1 mg/mL), compared with control (CTR). Quantitative parameters include: (A) number of master junctions; (B) total master segment length; (C) total branching length; (D) total segment length; (E) total mesh area; and (F) number of meshes. Data are expressed as mean ± SEM (n=3). Image analysis was performed using the Angiogenesis Analyzer plugin (ImageJ). Statistical analysis was performed using Student’s t-test. Asterisks indicate significance versus control: * p < 0.05 and ** p < 0.005.

#### 4.2.2 *In vitro* Inflammatory Model

##### 4.2.2.1 Effects of VFM on LPS-Induced Cell Viability

The exposure to LPS (25 µg/mL) significantly reduced pAECs viability after 24 hours (p < 0.01). The addition of VFM at high concentrations (1 mg/mL) to the culture system reduced LPS-induced cytotoxicity by restoring cell viability to basal levels; on the other hand, low concentrations of VFM (0.1 mg/mL) did not show to have any cytoprotective effect as cell viability was similar to cells only exposed to LPS and was significantly reduced compared to control cultures (p < 0.05) (Figure 6).

**Figure 6:**
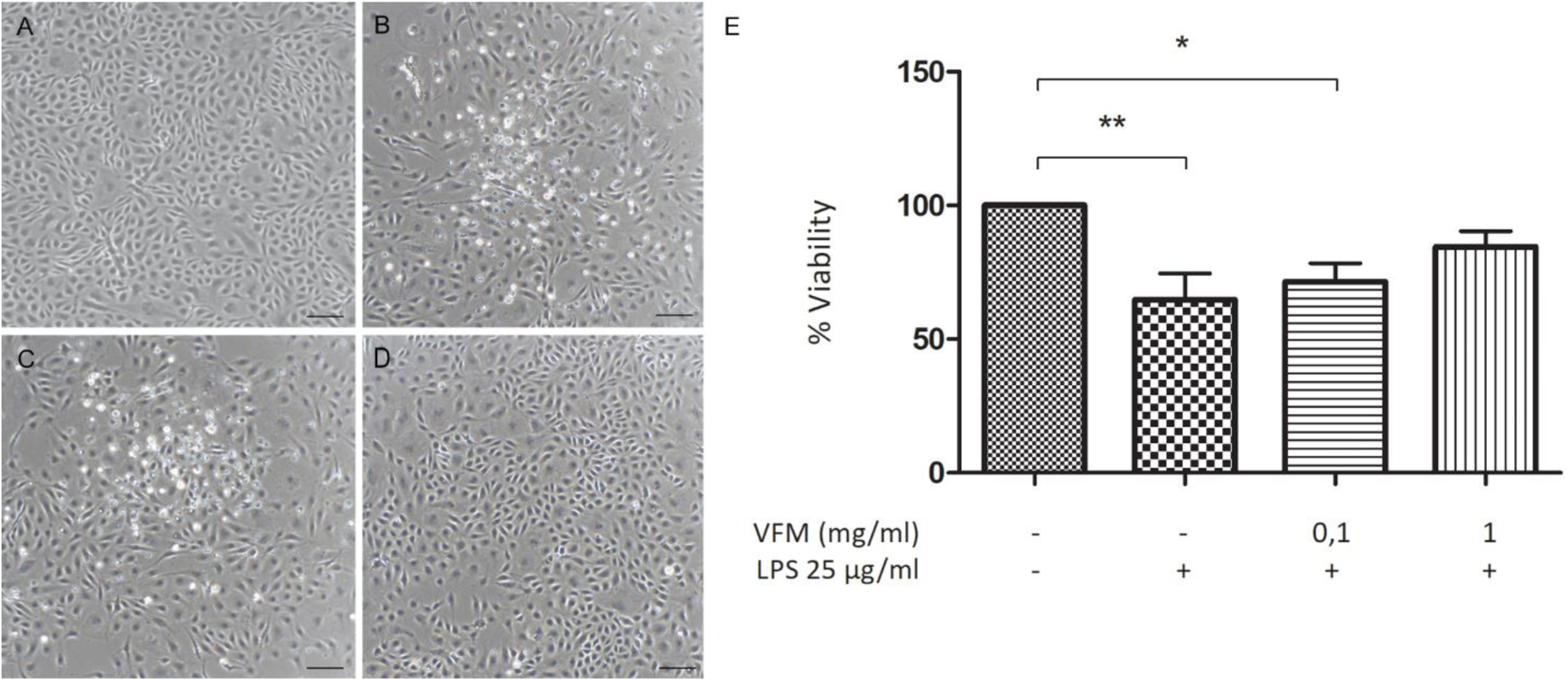
Effects of lipopolysaccharide (LPS) and Vaccinium floribundum Kunth microencapsulated extract (VFM) on cell viability as assessed by the MTT assay. Statistical analysis was performed using one-way ANOVA. *= p < 0.05; **=p < 0.01. Data are presented as mean ± SEM. n = 5.

##### 4.2.2.2 Effects of VFM on LPS-Induced Cytokine Gene Expression

IL-6 and IL-8 mRNA levels were significantly upregulated when pAECs were exposed to LPS alone compared to control conditions after 5 hours (Figure 7). The co-exposure of porcine endothelial cells to high doses of VFM (1 mg/mL) showed to partially counteract this LPS-induced response. Indeed, after 5 hours the concomitant addition of VFM with LPS led to a significant reduction of both pro-inflammatory cytokine gene expression compared to cells exposed to LPS alone; nonetheless, the mRNA levels were still significantly higher compared to control cells. Furthermore, the addition of VFM alone was able to significantly downregulate IL-8 gene expression levels compared to control cells (Figure 7).

**Figure 7:**
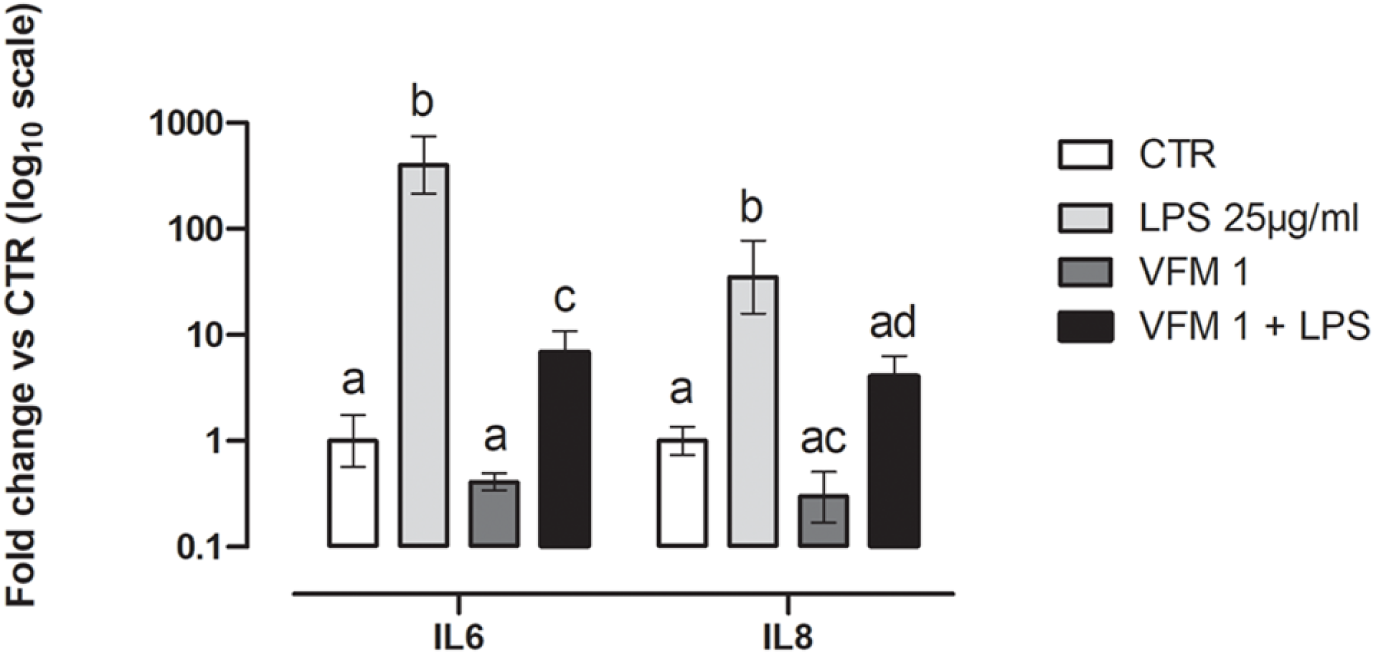
Effect of VFM on LPS-induced IL-6 and IL-8 gene expression. Relative mRNA levels of IL-6 and IL-8 were quantified by RT–qPCR at 5 h. Cells were treated with lipopolysaccharide (LPS) alone or in combination with VFM at the indicated concentration (1 mg/mL). Gene expression levels are reported as Fold Change relative to the control (2-ΔΔCt method) and presented on a log10 scale. Error bar represents the range of relative gene expression (n=3). Values not sharing a common letter indicate statistically significant differences (p < 0.05) among experimental groups, as determined by one-way ANOVA followed by post-hoc multiple comparisons. CTR, untreated control; LPS, lipopolysaccharide; VFM, Vaccinium floribundum Kunth microencapsulated extract.

### 4.3 *In vivo* model

#### 4.3.1 Inflammatory inhibition

To compare the different treatments, the percentage of inflammation inhibition was calculated. The results showed that the positive control (Diclofenac) had an average inhibition of 51.74±3.26%, while the microencapsulated group at 300 mg/kg had an average inhibition of 50.63±1.66%. There were no significant differences (p < 0.05), indicating that the anti-inflammatory effect of microencapsulation at this dose is comparable to that of the positive control treatment (Figure 8).

**Figure 8:**
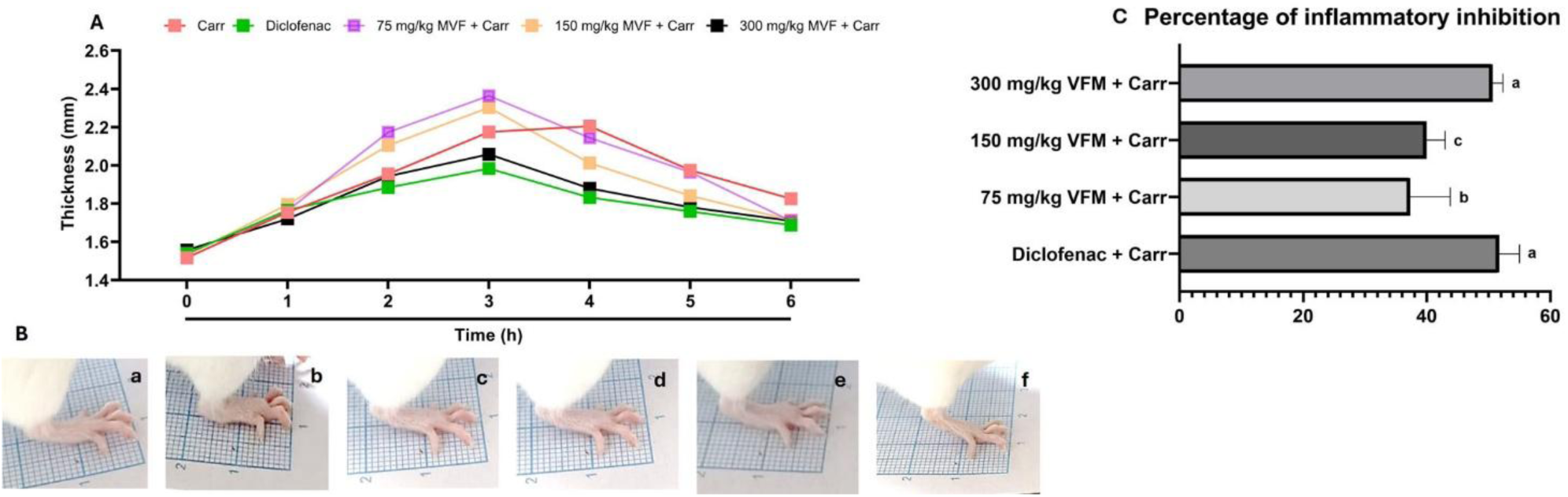
Effect of Vaccinium floribundum Kunth microencapsulated extract (VFM) in mice (n=5) by carrageenan paw edema assay. The thickness of paw (mm) was measured at 1, 2, 3, 4, 5 and 6 h after the injection of Carr respectively (A). Effects of VFM on the Inflammatory response at 6 h (B), control group (a), Carr group (b), Diclofenac group + Carr (c), 75 mg/kg VFM + Carr (d), 150 mg/kg VFM + Carr (e), 300 mg/kg VFM + Carr (f). Percentage of inflammatory inhibition (C). Statistical analysis was performed using one-way ANOVA and Dunnet post hoc-test. Data are presented as mean ± SD. n = 5

## 5 Discussion

For centuries, the local communities of Ecuador have leveraged the country’s vast floral biodiversity for traditional medicinal applications. Within this ethnobotanical landscape, berries of *Vaccinium* spp. have gained global recognition as "superfruits," due to their antioxidant capacity and potential health benefits (29). The synergy between historical traditional use, cultural heritage, and the identification of potent bioactive constituents, such as polyphenols and anthocyanins, has catalyzed significant interest in the pharmacological and nutritional exploration of these species (30). For this reason, this study was designed to characterize the phytochemical profile and biological activity of a microencapsulated extract of *Vaccinium floribundum* Kunth (VFM), a species central to Ecuadorian traditional medicine. Specifically, we investigated the biological effects of VFM on primary porcine aortic endothelial cells (pAECs), focusing on its cytocompatibility, its role in promoting angiogenesis through the modulation of key molecular receptors, and its protective efficacy against lipopolysaccharide (LPS)-induced endothelial dysfunction and inflammation. Furthermore, its anti-inflammatory potential was tested using an *in vivo* paw edema model which is a standardized acute, non-infectious model for inflammation. Microencapsulation represents an advanced delivery strategy designed to enhance the physicochemical stability of bioactive compounds by forming a protective matrix that limits environmental exposure (31). This approach is particularly relevant for phenolic compounds,which hydroxyl groups and conjugated structures are highly susceptible to photodegradation, thermal instability, and oxidative stress. Thus, the protective barrier former mitigates the formation of reactive oxygen species and preserves the intrinsic antioxidant and anti-inflammatory properties of the encapsulated compounds (19).

The microencapsulation process resulted in a process yield consistent with previously reported values for polysaccharide-based delivery systems, suggesting high encapsulation efficiency and limited material loss (32). FTIR analysis revealed spectral features characteristic of polyphenol–polysaccharide interactions, including enhanced absorption bands within the 1050–1160 cm⁻¹ region attributed to C–O–C glycosidic linkages and C–O stretching vibrations. The increased intensity and structural reinforcement of these bands further supported the stabilizing contribution of maltodextrin as a carrier matrix(33).

The phenolics profile observed in VFM was characterized by the dominance of B-type procyanidins, quercetin-based flavonols, and chlorogenic acid. The high concentration of Procyanidin B1 relative to B2 and C1 suggested a biosynthetic preference for specific dimeric structures, which are critical for the fruit’s astringency and the antioxidant capacity (34). The lower levels of procyanidin C1 (a trimer) compared to the B1 dimer reflected the typical oligomerization gradient where dimers are more prevalent than higher-order trimers (35). The flavonol profile was dominated by quercetin and its galactoside derivative. Blueberries are well-documented sources of quercetin, a potent anti-inflammatory agent, typically containing about 100 mg/kg (36). The presence of hyperoside (quercetin-3-*O*-galactoside) aligned with the plant’s tendency to store flavonols in glycosylated forms to increase stability and solubility (37). Likewise, the higher levels of kaempferol-7-*O*-glucoside compared to its aglycone reflected this metabolic storage strategy. The detection of phloretin and phloridzin was significant as these dihydrochalcones were more famously associated with apples, yet they were increasingly identified in blueberries as minor but potent bioactive markers (38). Furthermore, the levels of epicatechin provided the necessary substrate for the condensed tannins (procyanidins) identified, indicating a highly active flavonoid pathway (39). The dominance of chlorogenic acid aligned with its role as the hallmark antioxidant in blueberries, often constituting the majority of the total phenolic acid fraction (40). Its high concentration relative to its precursor, caffeic acid, indicated a high rate of esterification with quinic acid within the fruit’s metabolic pathways (20). The uniform, low levels of ferulic and *p*-coumaric acids suggested that these compounds served primarily as transient intermediates in the biosynthesis of more complex polyphenols or were integrated into the cell wall matrix (41). Meanwhile, the detection of protocatechuic acid was noteworthy as it often emerged from the degradation of anthocyanins or existed as a biosynthetic precursor to gallic acid derivatives (42).

The preliminary safety screening confirmed that VFM is highly cytocompatible with porcine aortic endothelial cells (pAECs). In fact, treatment of pAECs with VFM for 24 h did not negatively affect cell viability at any of the concentrations tested. Most importantly, at 1 mg/mL, VFM significantly enhanced cell metabolic activity. Therefore, the increased cell viability might suggest that VFM may actively support endothelial health and metabolic vigor. The observed bioactivity of VFM is likely due to the high content of polyphenols and anthocyanins found in *Vaccinium* species, which have been shown to modulate metabolic pathways and reduce oxidative stress in vascular models (43,44).

Following the preliminary assay, pAECs’ angiogenesis potential was evaluated using an *in vitro* extracellular matrix-based assay widely recognized for evaluating the ability of endothelial cells to organize into complex structures. The formation of new vascular networks is a multi-step process that requires the transition of endothelial cells from a quiescent state to an active, migratory, and organizational phenotype.

Our results demonstrate that VFM treatment significantly promotes the organizational capacity of pAECs. This is evidenced by a substantial increase in all evaluated morphometric parameters, including master junctions, total segment length, and mesh area (Fig. 5). The fact that the extract enhances the complexity of the capillary-like network across all metrics suggests that VFM does not merely stimulate random migration but actively orchestrates the cellular interactions necessary for structural stabilization. Interestingly, this morphological transformation was accompanied by a specific and transient transcriptional response. At the 5-hour mark, a significant up-regulation was observed exclusively for the FLT-1 (VEGFR-1) receptor. By 24 hours, the mRNA levels of all tested markers, including VEGF, FLK-1 (VEGFR-2), and HO-1, returned to baseline. In vascular biology, FLT-1 and FLK-1 are considered "early molecular switches" that trigger the endothelial-to-angiogenic transition. Given that FLT-1 plays a crucial role in guidance and spatial organization during the initial phases of vessel assembly. However, the return to baseline levels for all genes by 24 hours suggests that the 5-hour time point may represent the tail end of this initial transcriptional wave. Consequently, the observed lack of significant modulation at 24 hours, and the partial response at 5 hours, indicates that VFM initiates a rapid signalling cascade. Future studies should focus on earlier time points, such as 3 hours, to capture the peak expression of this “angiogenic switch” and fully characterize the kinetics of VFM-mediated endothelial activation.

Overall our results concerning angiogenesis are in agreement with the proangiogenic potential of polyphenols and their ability to orchestrate complex molecular signalling pathways that promoted the growth and stabilization of vascular networks (45). These natural compounds have been reported to regulate various stages of angiogenesis, including the modulation of factors such as basic fibroblast growth factor (bFGF: (46)), vascular endothelial growth factor (VEGF: (47,48)), hypoxia-inducible factor-1α (HIF-1α: (49)), and matrix metalloproteinase (MMP) activity (50). Moreover, they also influence endothelial cell functions, including proliferation, migration, and tube formation (51) and references therein, thereby impacting tumor vascularization and metastatic potential (52), as well as vascular-associated complications. Several studies carried out both *in vivo* and *in vitro* have demonstrated that angiogenesis represent a therapeutic target for plant polyphenols (51). These compounds, specifically at low doses, activated critical receptors such as VEGFR, EGFR, and the ANG 1/2/tyrosine kinase system, essential for endothelial cell proliferation and migration (53). Among polyphenol compounds detected in VFM, delphinidin is primarily recognized for its anti-angiogenic properties in cancer contexts, it could exhibit a pro-angiogenic or vasculoprotective effect under specific physiological conditions, often characterized by a biphasic (dose-dependent) response. At low concentrations, delphinidin and similar anthocyanidins could promote the release of nitric oxide (NO) from endothelial cells (54), enhanced blood flow and might support healthy vessel maintenance or repair in ischemic tissues through the activation of estrogen receptor alpha (ER*α*) (55). Essentially, it acted as a modulator that suppressed abnormal vessel growth while potentially supporting the integrity of existing healthy vasculature by supporting basal mitochondrial DNA and mRNA expression of factors like NRF1 and Tfam in normal cells (56).

Lipopolysaccharide (LPS) is a well-established inducer of endothelial dysfunction, triggering an inflammatory response along with oxidative stress, resulting in high cell death rate due to the cytotoxic environment (57). In the present study, LPS exposure significantly reduced endothelial cells viability, which might confirm the presence of an inflammatory microenvironment. Notably, high doses of VFM (1 mg/mL) restored cell viability almost to basal levels after 24 hours. This cytoprotective effect could be ascertained to the high concentrations of flavonoids and anthocyanins contained in VFM, which has been identified in the herein described extract (58–60). Indeed, the cytoprotection provided by mortiño extracts might be also reflected by the gene expression of pro-inflammatory interleukins mRNA levels (IL-6 and IL-8) as high doses of VFM significantly down-regulated their mRNA levels compared to pAECs exposed only to LPS after 5 hours of exposure. Different studies have demonstrated that blueberries extract, due to presence of compounds such as polyphenols, are able to modulate inflammatory and oxidative environments in complex biological systems (44,61,62). As a matter of fact, the bioactive constituents present in the mortiño fruit extract, namely flavonoids and anthocyanins, are known for being able to modulate the protective endogenous oxidative stress machinery of cells by upregulating the gene expression of nuclear factor erythroid 2-like 2 (Nfr2) (63) or by inhibiting NF-κB signaling pathway in endothelial cells (64–66) or blocking the same pathway along with the AMPK signalling one, which is a known mechanism of action of quercetin flavonols (67–69). These mechanisms together might have led to the observed reduction of inflammation and oxidative stress within LPS-stimulated pAECs, which showed a downregulated expression of both IL-6 and IL-8 after 5 hours in culture, along with the increased pAECs viability after exposure to LPS compared to LPS alone.

Finally, the results concerning *in vivo* model of acute inflammation by using the carrageenan-induced paw edema assay (70–72) demonstrated the anti-inflammatory property of VFM. In particular, following carrageenan administration, a rapid inflammatory response occured, characterized by edema, hyperalgesia, and erythema due to the release of several pro-inflammatory mediators including histamine, complement components, and pro-inflammatory cytokines (73). In the present study, all tested concentrations of the extract exhibited anti-inflammatory activity, with the highest dose (300 mg/kg) producing the most pronounced reduction in paw edema. These findings are consistent with previous reports describing anti-inflammatory properties in other species of the Vaccinium genus. For instance, reductions in carrageenan-induced inflammation of approximately 28.8% for V. myrtillus (200 mg/kg), 36.9% for V. macrocarpon (200 mg/kg), and 45.3% for V. myrtillus (200 mg/kg) have been reported at 5 h post-induction. Although these values are slightly lower than those observed in the present study, this difference may be explained by the microencapsulation of the extract, which could promote a more sustained release of bioactive compounds and prolong their anti-inflammatory activity throughout the experimental study (74).

In conclusion, the microencapsulated *Vaccinium floribundum* Kunth extract (VFM) demonstrated robust pro-angiogenic and anti-inflammatory activities in both *in vitro* and *in vivo* models. VFM effectively enhanced endothelial cell viability, promoted capillary-like network formation, and modulated early angiogenic signaling pathways, highlighting its potential to support vascular health. Moreover, VFM mitigated LPS-induced endothelial dysfunction by downregulating pro-inflammatory cytokines, consistent with the known bioactivity of polyphenols and anthocyanins. The *in vivo* paw edema assay further confirmed its anti-inflammatory efficacy, with microencapsulation likely contributing to sustained bioactive release. Altogether, these findings provide a scientific basis for the traditional use of *Vaccinium floribundum* in Ecuadorian ethnomedicine, highlighting the relevance of the traditional knowledge and supporting the sustainable use of natural resources in a country characterized by high biodiversity. Future perspectives may include the development of a VFM-based nutraceutical formulation targeting vascular and inflammatory disorders, although this will require overcoming the limitations usually associated with wild plant harvesting, ensuring the production of standardized and more chemically stable extract.

## 2 Author Contributions

FA, IT: Conceptualization, Funding acquisition, Supervision, Writing – original draft, Formal analysis, Writing – review & editing, Resources, Validation. GB, LM: Investigation, Methodology, Formal analysis, Data curation, Writing – original draft, Writing – review & editing. RDL, RS, CVG, GOZC: Methodology, Investigation, Data curation, Writing – review & editing. SMA: Methodology, Investigation, Data curation, Writing original draft– review & editing, Validation. GV, MF: Writing – review & editing, Investigation, AZ, Resources, Writing – review & editing, Funding acquisition, Formal analysis, Supervision, Data curation, Methodology, Investigation. CB: Conceptualization, Resources, Writing – review & editing, Funding acquisition, Formal analysis, Supervision, Data curation, Methodology, Investigation.

## 4 Acknowledgments

NA

## Data Availability Statement

The datasets *VFM-pAEC* for this study can be found in the AMSACTA from the University of Bologna https://amsacta.unibo.it/id/eprint/8873

